# The PASTA domains of *Bacillus subtilis* PBP2B stabilize the interaction of PBP2B with DivIB

**DOI:** 10.1101/713677

**Authors:** Danae Morales Angeles, Alicia Macia-Valero, Laura C. Bohorquez, Dirk-Jan Scheffers

## Abstract

Bacterial cell division is mediated by a protein complex known as the divisome. Many protein-protein interactions in the divisome have been characterized. In this report, we analyse the role of the PASTA (Penicillin binding protein And Serine Threonine kinase Associated)-domains of *Bacillus subtilis* PBP2B. PBP2B itself is essential and cannot be deleted, but removing the PBP2B PASTA domains results in impaired cell division and a heat sensitive phenotype. This resembles the deletion of *divIB*, a known interaction partner of PBP2B. Bacterial two hybrid and co-immunoprecipitation analyses show that the interaction between PBP2B and DivIB is weakened when the PBP2B PASTA domains are removed. Combined, our results show that the PBP2B PASTA domains are required to stabilize the interaction between PBP2B and DivIB.

## Introduction

The synthesis of peptidoglycan during cell division is essential for the completion of division and in fact considered one of the drivers for constriction itself (1, 2). Cell division is mediated by a complex of proteins collectively known as the divisome. In most bacteria, the divisome contains two division specific peptidoglycan synthesis proteins, FtsW, a protein from the SEDS-family with glycosyl transferase activity, and a division specific Class B Penicillin Binding Protein (bPBP) with transpeptidase activity (3, 4). In *Bacillus subtilis*, these proteins are FtsW and PBP2B, which are both essential (5, 6). Interestingly, recent work from our lab and of Daniel and colleagues has shown that it is the presence of PBP2B that is essential, rather than its transpeptidase activity, suggesting that the function of PBP2B is a scaffolding one (7, 8). This was similar to a previous report on the *Streptococcus pneumoniae* homologue PBP2x, of which the transpeptidase activity is also not essential (9). Both PBPs contain PASTA domains at their C-terminus. PASTA – for Penicillin binding protein And Serine Threonine kinase Associated – domains are exclusively found in Gram-positive bacteria, in some high molecular weight PBPs and in eukaryotic-like serine/threonine kinases (eSTKs) (10). These domains, which contain 60-70 amino acids, have a characteristic secondary structure which consists of three β strands and a α helix; the first and the second β strands are connected by a loop, but the sequence of the domain is not well conserved. PASTA domains can be present as a single or multiple copies in proteins. In PBP2x, loss of the PASTA domains abolishes the binding of Bocillin-FL, a fluorescent penicillin derivative (11) and localization of PBP2x to the division site (9), suggesting that PASTA domains mediate the interaction with peptidoglycan. This was nicely illustrated recently by a series of crystal structures that revealed that the PBP2x PASTA domains form an allosteric binding site for a pentapeptide stem in a nascent peptidoglycan strand, which positions another peptide stem on the same strand in the active site so that it can be cross-linked (12). The allosteric binding site is formed at the interface of the two PASTA domains and the transpeptidase domain and comprises the entire first and part of the second PASTA domain. Binding of the terminal D-Ala-D-Ala of the stempeptide at this side displaces a ‘gatekeeper’ Arginine residue on the transpeptidase domain, which subsequently forms salt bridges with an Aspartate and a Glutamate residue on the first PASTA domain which opens up the active site so that the donor stempeptide for transpeptidation on the same glycan strand can bind (12). The PASTA domains of *B. subtilis* PBP2B lack all the residues required for this allosteric activation.

Not all PASTA-domain containing proteins bind peptidoglycan (13). Bioinformatics analyses revealed a key difference between PASTA domains that bind peptidoglycan and PASTA domains from proteins that don’t – in a residue that determines the flexibility of the “putative binding pocket”, a conserved region localized at the end of the β strand. Binder PASTA domains have an Arginine or a Glutamate residue at this position, while non-binders have a Proline (13). An example of this is *B. subtilis* PrkC, which functions as a peptidoglycan fragment sensor that induces spore germination (14), in which mutation of this Arginine abolishes peptidoglycan binding (15). PBP2B has Prolines at both sites in its PASTA domains. Thus, PBP2B does not have residues associated with peptidoglycan binding by its individual PASTA domains nor with an allosteric site formed between the PASTA domain and the transpeptidase domain. Combined with our previous observation that deletion of the PBP2B PASTA domains does not affect localization or binding of Bocillin-FL (7) this suggests that the PASTA domains of PBP2B have a different function than peptidoglycan binding.

Other reported functions for PASTA domains include protein localization and kinase activation (16). In *S. pneumoniae* StkP, which contains 4 PASTA domains, the 4^th^ domain is critical for localization through interaction with the peptidoglycan hydrolase LytB, whereas the first three PASTA domains function as a ruler that positions the 4^th^ domain to control cell wall thickness (17).

In this paper, we have further investigated the role of the PASTA domains of PBP2B and show that these domains strengthen the interaction between PBP2B and the divisome protein DivIB. This interaction becomes critical when cells are grown at higher temperatures.

## Methods

### Strains and media

Strains used in this study are listed in Table 1. All *Bacillus* strains were grown in casein hydrolysate (CH)-medium at 30°C unless other conditions are specified. When necessary kanamycin (5 µg/ml) and spectinomycin (50 µg/ml) were added. To induce the expression of genes under control of the P_spac_ and P_xyl_ promoters, either isopropyl β-D-1-thiogalactopyranoside (IPTG) (0.5 mM) or xylose (0.2% w/v) were added to the medium.

**Table 1.**
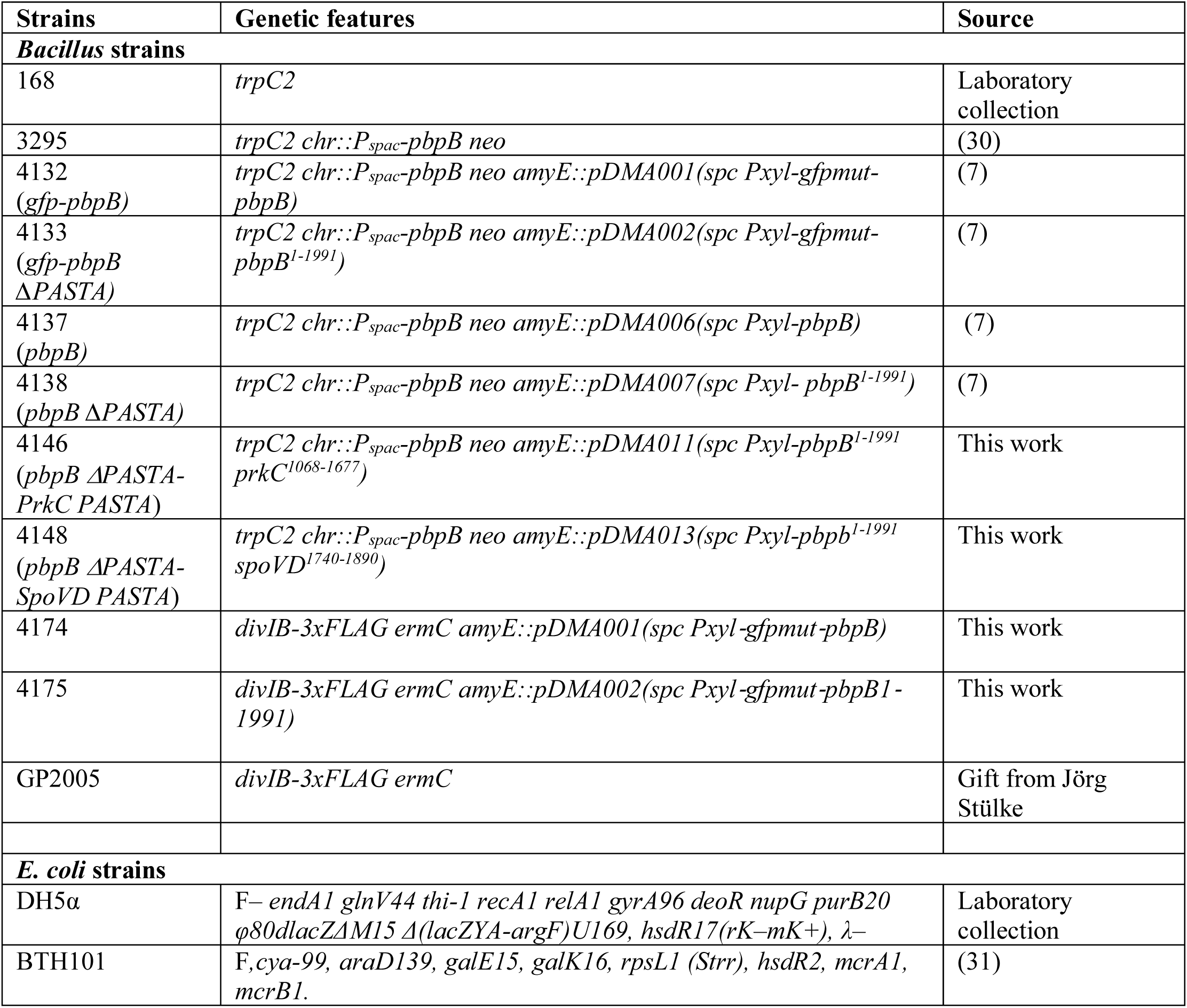
Strains.

| Strains | Genetic features | Source |
| --- | --- | --- |
| ***Bacillus* strains** |  |  |
| 168 | *trpC2* | Laboratory collection |
| 3295 | *trpC2 chr::P<sub>spac</sub>-pbpB neo* | (30) |
| 4132<br>(*gfp-pbpB*) | *trpC2 chr::P<sub>spac</sub>-pbpB neo amyE::pDMA001(spc Pxyl-gfpmut-pbpB)* | (7) |
| 4133<br>(*gfp-pbpB ΔPASTA*) | *trpC2 chr::P<sub>spac</sub>-pbpB neo amyE::pDMA002(spc Pxyl-gfpmut-pbpB<sup>1-1991</sup>)* | (7) |
| 4137<br>(*pbpB*) | *trpC2 chr::P<sub>spac</sub>-pbpB neo amyE::pDMA006(spc Pxyl-pbpB)* | (7) |
| 4138<br>(*pbpB ΔPASTA*) | *trpC2 chr::P<sub>spac</sub>-pbpB neo amyE::pDMA007(spc Pxyl- pbpB<sup>1-1991</sup>)* | (7) |
| 4146<br>(*pbpB ΔPASTA-PrkC PASTA*) | *trpC2 chr::P<sub>spac</sub>-pbpB neo amyE::pDMA011(spc Pxyl-pbpB<sup>1-1991</sup> prkC<sup>1068-1677</sup>)* | This work |
| 4148<br>(*pbpB ΔPASTA-SpoVD PASTA*) | *trpC2 chr::P<sub>spac</sub>-pbpB neo amyE::pDMA013(spc Pxyl-pbpB<sup>1-1991</sup> spoVD<sup>1740-1890</sup>)* | This work |
| 4174 | *divIB-3xFLAG ermC amyE::pDMA001(spc Pxyl-gfpmut-pbpB)* | This work |
| 4175 | *divIB-3xFLAG ermC amyE::pDMA002(spc Pxyl-gfpmut-pbpB<sup>1-1991</sup>)* | This work |
| GP2005 | *divIB-3xFLAG ermC* | Gift from Jörg Stülke |
| ***E. coli* strains** |  |  |
| DH5α | F– *endA1 glnV44 thi-1 recA1 relA1 gyrA96 deoR nupG purB20 φ80dlacZΔM15 Δ(lacZYA-argF)U169, hsdR17(rK–mK+), λ–* | Laboratory collection |
| BTH101 | F, *cya-99, araD139, galE15, galK16, rpsL1 (Strr), hsdR2, mcrA1, mcrB1*. | (31) |

### Construction of PBP2B chimeras

Chimeras (Figure S1) were constructed using restriction free cloning (18). Hybrid primers were used to amplify *prkC* and *spoVD* regions coding for PASTA domains from chromosomal DNA of *B. subtilis*. The hybrid primers were designed using http://www.rf-cloning.org/, primers (Table S1) contain complementary sequences to *prkC* or *spoVD* and plasmids pDMA002 or pDMA007. A first PCR was performed using the hybrid primers to create a mega-primer which contains *prkC* or *spoVD* PASTA domains flanked by complementary sequences of pDMA002 or pDMA007. The mega-primers were used in a second PCR to replace the *pbpB* PASTA domains from pDMA002 or pDMA007 with *prkC* or *spoVD* PASTA domains. *Dpn*I was added to the products obtained in the second PCR in order to degrade the original plasmid. After digestion, the PCR products were used to transform *E. coli* DH5α cells. Resulting plasmids (Table 2) were sequenced and cloned into *amyE* locus of *B. subtilis* 3295. Integration into the *amyE* locus was verified by growing the transformants on starch plates.

**Table 2.** Plasmids.

| Plasmid | Genetic features | Source |
| --- | --- | --- |
| pDMA002 | *bla amyE3' spc Pxyl- gfpmut1-pbpB<sup>1-1991</sup> - ' amyE5'* | (7) |
| pDMA007 | *bla amyE3' spc Pxyl- pbpB<sup>1-1991</sup> amyE5'* | (7) |
| pDMA011 | *bla amyE3' spc Pxyl- pbpB<sup>1-1991</sup> prkC<sup>1068-1677</sup>' amyE5'* | This work |
| pDMA012 | *bla amyE3' spc Pxyl-gfpmut1 pbpB<sup>1-1991</sup> prkC<sup>1068-1677</sup>' amyE5'* | This work |
| pDMA013 | *bla amyE3' spc Pxyl- pbpB<sup>1-1991</sup> spoVD<sup>1735-1916</sup>' amyE5'* | This work |
| pDMA014 | *bla amyE3' spc P<sub>xyl</sub>-gfpmut1- pbpB<sup>1-1991</sup> spoVD<sup>1735-1916</sup>)' amyE5'* | This work |
| pKT25 | Plasmid encoding T25 fragment of *B. pertussis* *cyaA*; Km<sup>R</sup> | (31) |
| pUT18C | Modified version of pUT18 with the polylinker located on the C-terminal end of T18; Amp<sup>R</sup> | (31) |
| pKT25-zip | Derivative of pKT25 with a leucine zipper of GCN4 fused to the T25 fragment, Km<sup>R</sup> | (31) |
| pUT18C-zip | Derivative of pUT18C with leucine zipper of GCN4 fused to the T18 fragment, Amp<sup>R</sup> | (31) |

### Growth curves

Strains were grown overnight in the presence of kanamycin (5 µg/ml) and spectinomycin (50 µg/ml) when necessary. IPTG was added to the medium to express wild-type *pbpB* and to ensure the proper growth of all strains before performing the growth curves. The following day, the strains were diluted to an OD_600_ 0.05 and grown until early exponential phase. Next, cells were washed CH-medium to remove the IPTG. Cells were diluted to an OD_600_ 0.001 in CH-medium containing 0.2% (w/v) xylose to express PBP2B, PBP2B-ΔPASTA or PBP2B chimeras. 200 µl of culture (in triplicate), of each condition to test, was loaded in a 96-well plate. The cultures were grown at 30 or 48 °C, OD_600_ was measured every 10 minutes and recorded using a Powerwave 340 (Biotek).

### Microscopy

Cells were grown until exponential phase. Nile red (Sigma) (5µg/ml) and 4’,6-diamidino-2-phenylindole (Sigma) (DAPI) (1 µg/ml) were used to stain membranes and DNA, respectively. Cells were spotted on agarose (1% w/v in PBS) pads and imaged using a Nikon Ti-E microscope (Nikon Instruments, Tokyo, Japan) equipped with a Hamamatsu Orca Flash4.0 camera. Image analysis was performed using the software packages ImageJ (http://rsb.info.nih.gov/ij/), ObjectJ (https://sils.fnwi.uva.nl/bcb/objectj/index.html), ChainTracer (19) and Adobe Photoshop (Adobe Systems Inc., San Jose, CA, USA). Box plots were generated using BoxPlotR (http://shiny.chemgrid.org/boxplotr/). All quantitative results were derived from at least two biological replicate experiments.

### Protein stability

Membranes from strains 4132 and 4133 grown at 30°C on CH with 0.2% (w/v) xylose were isolated. Cells were grown until exponential phase and spun down (3 000 rpm, 7 min, 4 °C). Pellets were washed in PBS and then cells were lysed by sonication. Membranes were collected by centrifugation (45,000 rpm, 4°C, 50 min) and resuspended in PBS. The protein concentration was equalised for the two strain samples and aliquots of membrane material of the same volume were prepared. Aliquots were incubated at 30 or 48 °C for 5 min, 20 min, 1 hr, 2 hr and 14 hr. Then, Bocillin 650/665 (5 µg/ml) was added to each sample, and samples were further incubated at RT for 10 min. After incubation, sample buffer was added to each sample to stop further protein degradation, and samples were run in SDS (10 %) gel. GFP and Bocillin were detected using a Typhoon FLA950 (GE Healthcare). For GFP, the 473 nm laser and the LPB (Long Pass Blue) filter were used, and for Bocillin the 635 nm laser and the LPR (Long Pass Red) were used.

After imaging, the same gels were used for immunoblotting. Proteins were transferred to a PVDF membrane. Primary antibodies were anti-GFP (Thermofisher). Anti-Rabbit IgG alkaline phosphatase conjugated secondary antibodies (Sigma Aldrich) were used. Blots were developed using CDP-Star (Roche) and chemiluminescence was detected using a Fujifilm LAS 4000 imager (GE Healthcare).

### Bacterial two hybrids

Bacterial two hybrids were performed using the BACTH system components (kindly provided by Fabian Commichau, Göttingen University). Sequences from *divIB, divIC, ftsL, pbpb* and *pbpbΔPASTA* were amplified from chromosomal DNA of *B. subtilis* 168. Primers contained *Xba*I and *Kpn*I restriction sites (Table S1). Fragments were cloned into pKT25 and pUT18C using *Xba*I and *Kpn*I. The resulting plasmids were sequence verified and co-transformed into *E. coli* BTH101. To test for protein interactions, the transformants were plated on LB agar plates containing X-gal (40 µg/ml), IPTG (0.5 mM), kanamycin (50 µg/ml) and ampicillin (100 µg/ml). Plates were incubated at 30°C for 36 hrs and scored for blue color development. The β-Galactosidase assay was performed as described (20) with some modification. *E. coli* BTH101 containing the plasmids to test were grown as overnight cultures in LB containing IPTG (0.5 mM), kanamycin (50 µg/ml) and ampicillin (100 µg/ml) at 30°C. The next day 200 µl of cells were transferred to a tube containing buffer Z. To permeabilize the cells 20 µl of 0.01% SDS (w/v) and 40 µl of chloroform were added to each tube. After mixing, the chloroform was allowed to settle down and 50 µl of permeabilized cells were transferred to a 96-well plate containing 150 µl of buffer Z. Then, 40 µl of 4% (w/v) o-nitrophenyl-β-D-galactopyranoside (ONPG) was added to start the enzymatic reaction. When the samples were yellow, the reaction time was recorded and reactions were stopped by adding 1M Na_2_CO_3_ (final concentration). The absorbance at 420 nm and 550 nm was measured in a Powerwave 340 (Biotek) and β-Galactosidase activity was calculated as:

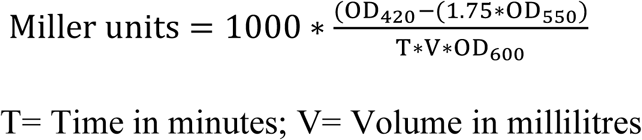

### Co-Immunoprecipitation

O/N cultures of strains 4174 and 4175 were diluted 1:100 and induced with 0.2% xylose (w/v) until OD_600_ ≈ 0.4. CoIPs were performed essentially as described (21). Cells were harvested, resuspended in buffer I (10 mM Tris-HCl, 150 mM NaCl, pH 7.4 with cOmplete™ ULTRA Tablets Mini EDTA-free, EASYpack protease inhibitors (Sigma-Aldrich)) and disrupted via sonication. Cell debris were removed by low-speed centrifugation and membranes were isolated through ultracentrifugation (100,000 × g, 1 h, 4°C) and solubilised with 1% (w/v) *n*-dodecyl-β-d-maltopyranoside (DDM; Anatrace) in buffer I by gentle shaking (4°C, 30 min). Solubilised material was recovered as the supernatant from a second ultracentrifugation step (100,000 × g, 30 min, 4°C). The protein concentration was determined with the DC™ (detergent compatible) protein assay kit (Bio-Rad Laboratories) and 200 ng total membrane proteins were incubated for 1 h at 4°C with gentle shaking on a roller mix with either 25 µl GFP-Trap® agarose beads (Chromotek) in a final volume of 100 µl 1% (w/v) DDM buffer I, according to manufacturers’ recommendations. Beads had been previously blocked by 1 h incubation with 1% (w/v) BSA in the corresponding buffer. After incubation, the flow-through fraction was collected (100 µl) using centrifugation (2,500 × g for 2 min) at 4°C and beads were washed twice and resuspended in 40 µl of 1xSDS-PAGE sample buffer. Low-binding tubes (Thermo Fisher Scientific) were used during the whole process. The input, flow-through and eluate fractions were analysed by SDS-PAGE and Western Blotting. Blots were developed using anti-FLAG M2 mouse monoclonal (Sigma-Aldrich, 1:1000) or anti-GFP pAb rabbit polyclonal (Chromotek, 1:1000) and appropriate Alkaline-phosphatase conjugated secondary antibodies. Blots were developed with CDP-Star (Roche), chemiluminescence was detected using a Fujifilm LAS4000 luminescence imager (GE Healthcare Life Science) and analysed using Image J (rsb.info.nih.gov/ij/).

## Results and Discussion

### The absence of PBP2B PASTA domains results in a temperature sensitive phenotype

In a previous study, we created a series of strains expressing PBP2B variants from which the PASTA domains were removed and/or the active site Serine was mutated (Figure S1). As *pbpB* is an essential gene, we generated strains in which the expression of wild type *pbpB* is under control of IPTG with an extra copy of the *pbpB* variant (with/without PASTA, with/without *gfp*) inserted in the *amyE* locus under control of the *P*_*xyl*_ promoter. This strategy allows cultivation of the strains while expressing wild type *pbpB* followed by depletion of PBP2B and a switch to PBP2B variant production by the removal of IPTG and the addition of xylose, thus ensuring that the observed phenotype is not a product of a suppressor mutation. Previously, we showed that PBP2B-ΔPASTA was able to complement PBP2B depletion under standard conditions (CH-medium, 30°C), indicating that PASTA domains are not essential under standard conditions (7). However, we noted that the cells were slightly elongated, which we have now quantified. The strain producing PBP2B has an average length of 3.34 µm (*n*= 200 cells), while the strain producing PBP2B-ΔPASTA has an average length of 4.85 µm (*n*= 200 cells), which is ∼1.5 times longer (Figure 1, Table S2). In addition, there is more variation in the length distribution of the PBP2B-ΔPASTA producing strain as can be observed in the boxplot. When the temperature was increased to 37°C, the average length of the PBP2B-ΔPASTA strain increased to 5.20 µm (*n=*200 cells), whereas the strain expressing PBP2B (*n=*200 cells) was slightly shorter than at 30°C (Figure 1, Table S2). We repeated this experiment in a defined minimal medium (SM medium) (*n*=200 per strain) and noticed a similar elongated phenotype for the strain expressing PBP2B-ΔPASTA compared to the strain expressing PBP2B (Figure S2). At 37°C, cells grown on SM medium were overall shorter, but again, the strain expressing PBP2B-ΔPASTA displayed an elongated phenotype.

**Figure 1.**
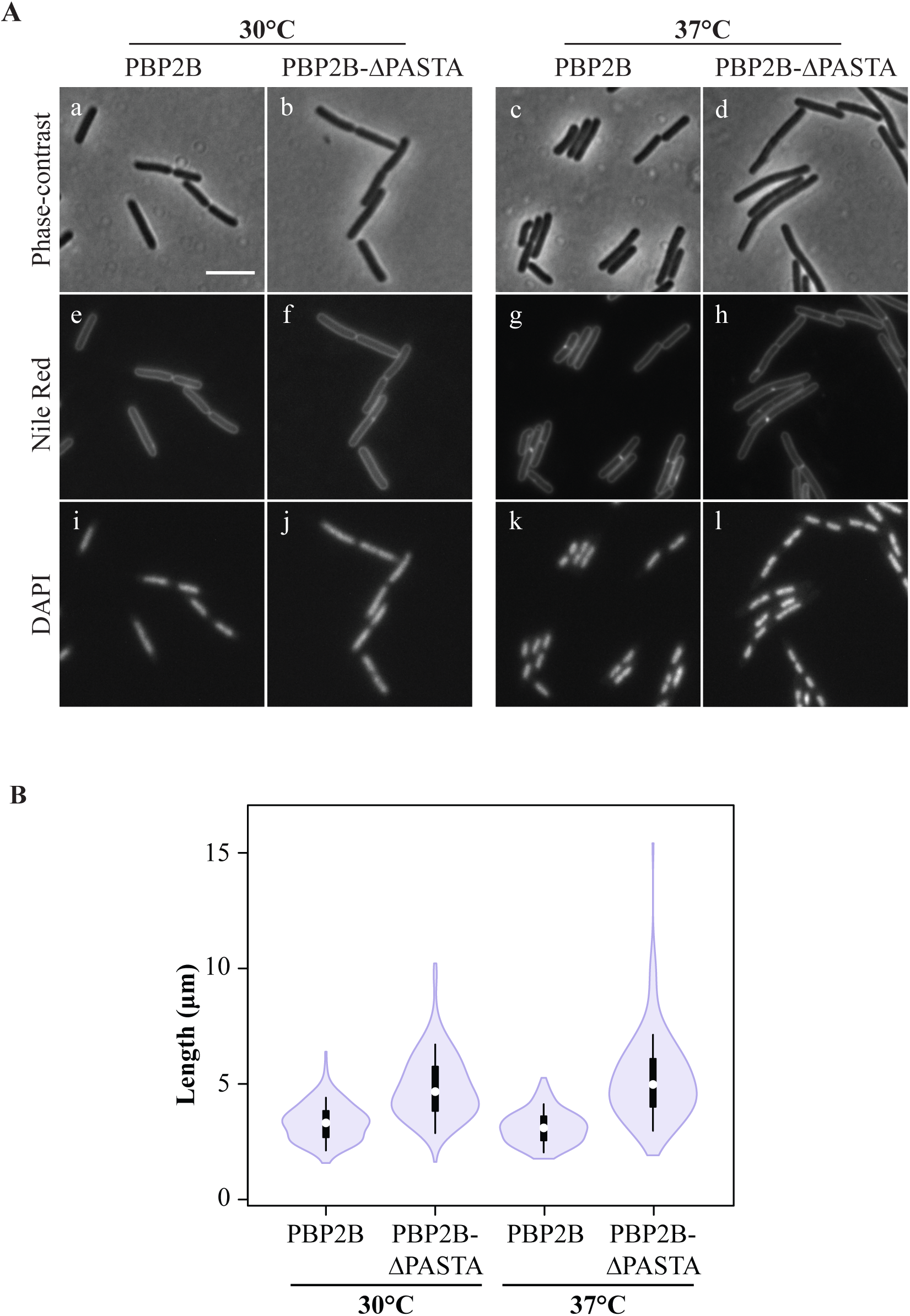
Phenotype of strains producing PBP2B and PBP2B-ΔPASTA. A) Phase-contrast microscopy of the PBP2B and PBP2B-ΔPASTA strains. Cultures were grown on CH-medium at 30°C and 37°C until exponential phase. Membranes and DNA were labelled with Nile red (e-h) and DAPI (i-l), respectively. (a, e, i) PBP2B (strain 4137) 30 °C; (b, f, j) PBP2B-ΔPASTA (strain 4138) 30°C; (c, g, k) PBP2B 37°C; (d, h, l) PBP2B-ΔPASTA 37°C. Scale bar: 5 µm, same for all. B) Length distribution of cells. Cells were grown in CH-medium at 30 or 37°C until exponential phase. As *B. subtilis* forms chains, cells were labelled with Nile red in order to determine the boundaries of single cells. Length of cells was obtained by automated image analysis. The values obtained (*n* = 200 per strain) are shown as box plots. White circles show the medians (PBP2B 30°C= 3.32 μm, PBP2B-ΔPASTA 30°C = 4.67 μm, PBP2B 37°C = 3.10 μm, PBP2B-PASTA 37°C = 4.97 μm); box limits indicate the 25th and 75th percentiles as determined by R software; whiskers extend 1.5 times the interquartile range from the 25th and 75th percentiles; polygons represent density estimates of data and extend to extreme values.

The increase in cell length is a characteristic phenotype indicative of a problem in cell division. To discard the possibility that the delay in cell division was a consequence of problems with chromosome segregation, DAPI was used to stain DNA. The PBP2B and PBP2B-ΔPASTA strains grown at 30°C and 37°C presented condensated nucleoids in all cells (Figure 1), indicating that chromosome segregation was not affected. Finally, strains expressing GFP-fusions to PBP2B and PBP2B-ΔPASTA, grown at 30°C, were scored for the presence of the GFP-PBP2B variant at the division site. GFP-PBP2B was present at the division site in 58.8% (± 4.9%, *n=* 609) of the cells, whereas GFP-PBP2B-ΔPASTA was present at the division site in 38.8% (±1.0%, *n*= 606) of the cells, again indicating that cell division is delayed/impaired when the PASTA domains are absent.

As we noticed that the elongation phenotype in CH-medium was more severe at 37°C than at 30°C, the temperature was increased to 48°C. Surprisingly, the PBP2B-ΔPASTA strain did not grow at 48°C (Figure 2A). This result suggests that the PBP2B-ΔPASTA strain is temperature sensitive. We also noted that after prolonged incubation, the control depletion strain 3295, started growing (Figure 2A) – this also happened at lower temperatures and analysis of several depletion strains revealed that this is due to appearance of suppressor mutations in the promoter used to control *pbpB* (not shown). To get more insight into the effects of high temperature on the phenotype, the strains were grown under normal conditions (30°C, CH-medium) to make sure that the cells were growing healthy. Then, cultures were shifted to 48°C and pictures were taken every 20 minutes. The strain expressing PBP2B showed no drastic changes in the phenotype during the course of the experiment (Figure 2B). On the other hand, after 40 minutes the PBP2B-ΔPASTA strain started to display cells with decreased contrast, a characteristic of dying cells (Figure 2B). After 1 hour at 48°C, we observed that the amount of dying cells in the PBP2B-ΔPASTA strain culture increased. These observations confirm that the deletion of the PASTA domains from PBP2B confers a temperature sensitive phenotype.

**Figure 2.**
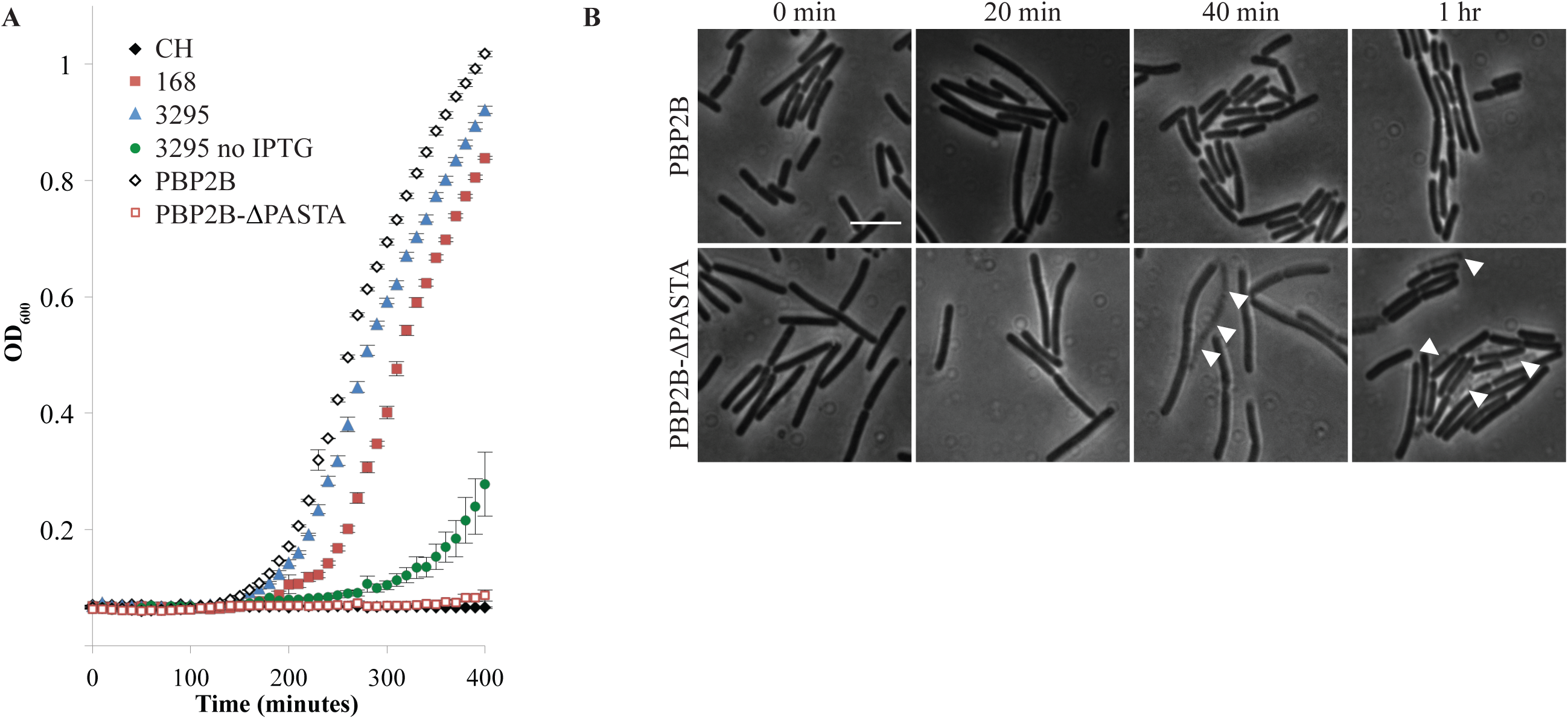
The PBP2B-ΔPASTA strain is thermosensitive. A) Growth curves in CH-medium at 48°C, OD_600_ was measured every 10 min. (♦) CH-medium, control, (▪)168, (▴) 3295, (•) 3295 no IPTG, (◊) PBP2B, (□) PBP2B-ΔPASTA. Representative results from three independent experiments are shown. All experiments were performed in triplicates. The resulting average and standard error are shown for each time point. B) Lysis of cells at 48°C. Cells were grown at 30°C until early exponential phase, then cells were shifted to 48°C and followed by microscope every 20 minutes. White arrowheads indicate dead cells. Scale bar 5 µm. Representative results from three independent experiments are shown.

### The PASTA domains of PBP2B are specific

*B. subtilis* has two other proteins that contain PASTA domains, SpoVD and PrkC. SpoVD is a PBP paralogous of PBP2B. It is crucial for spore cortex synthesis and contains a single PASTA domain (22). PrkC is a eukaryotic-like serine/threonine kinase that is involved in processes like germination and biofilm formation, WalR activation and that localizes to the septum (14, 23, 24). In order to test if the PASTA domains of SpoVD and PrkC were able to replace the function of the PASTA domains of PBP2B, the PBP2B PASTA domains were exchanged for PASTA domains from SpoVD or PrkC (Figure S1). Growth of the strains expressing the chimera proteins was followed at 30°C in CH-medium (Figure 3A), and was found to be similar to the background deletion strain. This indicates that the exchange of the PASTA domains did not interfere with the essential function of PBP2B. The cells expressing the chimera proteins were examined by microscopy which showed that the cells expressing the PBP2B-PASTA_SpoVD_ chimera were similar sized to cells expressing PBP2B, whereas cells expressing the PBP2B-PASTA_PrkC_ chimera were elongated, although not to the same extent as the PBP2B-ΔPASTA cells (Figure 3C, E, Table S2). GFP-fusions to the chimera proteins showed that both chimeras localize to division sites, as expected from the observation that the chimeras do not interfere with the essential function of PBP2B (Figure 3D). Subsequently, these strains were grown at 48°C to see whether the chimeric proteins complemented the temperature sensitive phenotype. Although both chimeric proteins did allow some growth at 48°C, the lag phase of the cells was longer compared to the PBP2B strain and cells did not reach similar OD_600_ values (Figure 3B). Again, the strain expressing the PBP2B-PASTA_PrkC_ chimera was most affected. These results indicate that the PASTA domains from other *B. subtilis* proteins can partially complement the absence of the PBP2B PASTA domains.

**Figure 3.**
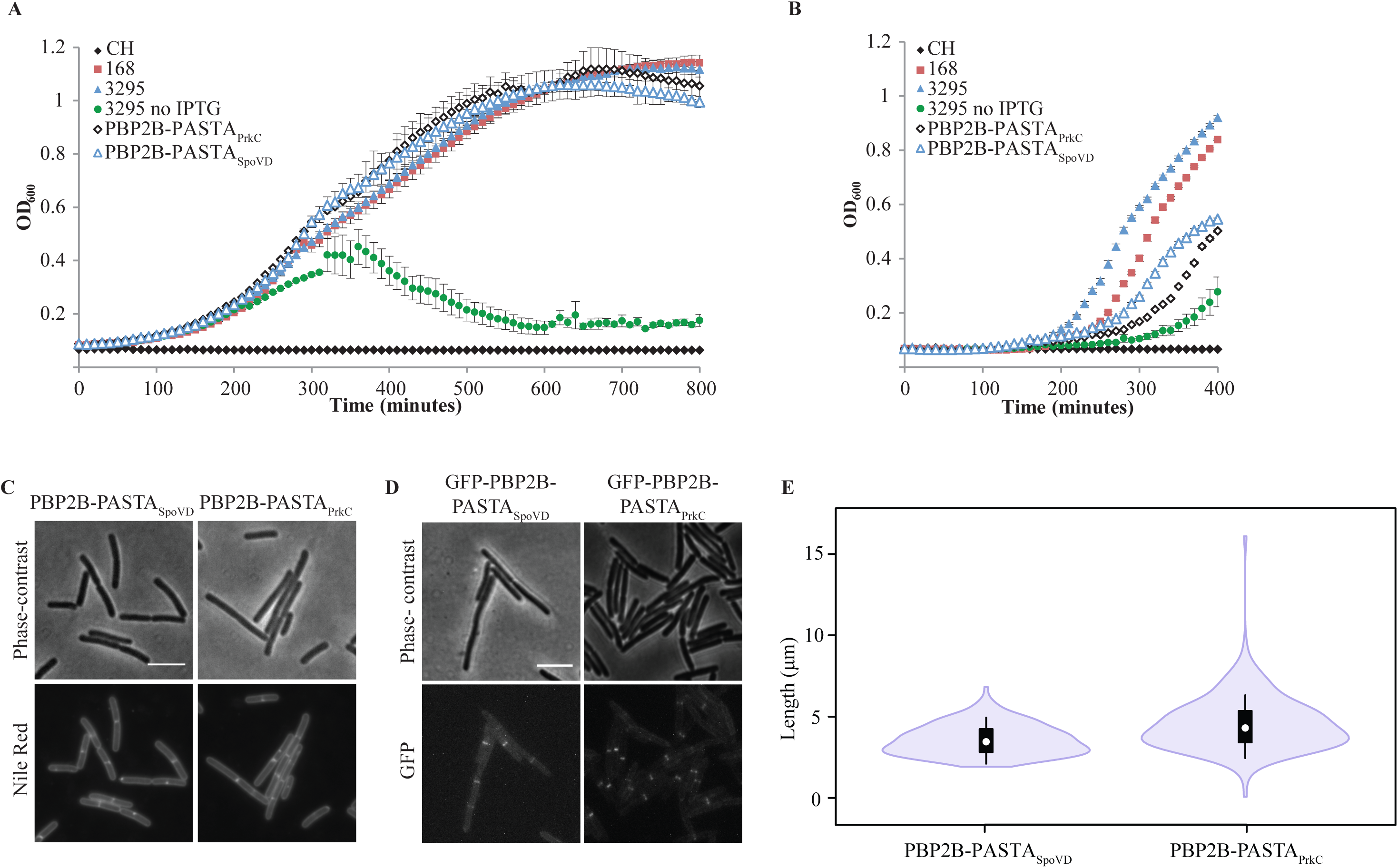
Phenotype of strains expressing PBP2B PASTA chimeras. Growth curves in CH-medium at 30°C (A) and at 48°C (B), OD_600_ was measured every 10 min. (♦) CH-medium, control, (▪)168, (▴) 3295, (•) 3295 no IPTG, (◊) PBP2B-PASTA_PrkC_, (Δ) PBP2B-PASTA_SpovD_. Representative results from three independent experiments are shown. All experiments were performed in triplicates. The resulting average and standard error are shown for each time point. C) Cells expressing PBPB2-PASTA_SpoVD_ and PBP2B-PASTA_PrkC_ were grown at 30°C in CH-medium until exponential phase, imaged with phase-contrast and labeled with Nile-red. Red. Representative results from three independent experiments are shown. Scale bar: 5 µm. D) Cells producing GPF-PBP2B-PASTA_SpovD_ and GFP-PBP2B-PASTA_PrkC_, imaged with phase-contrast and for GFP-fluorescence. Scale bar: 5 µm. E) Length distribution of cells producing PBP2B-PASTA_SpoVD_ and PBP2B-PASTA_PrkC_ chimeras. Cells were grown in CH-medium at 30°C until exponential phase and pictures were taken. Length of cells was obtained by automated image analysis. The values obtained (*n* = 200 per strain) are shown as box plots. White circles show the medians medians (PBP2B-SpoVD = 3.46 μm, PBP2B-PrkC=4.30 μm); box limits indicate the 25th and 75th percentiles as determined by R software; whiskers extend 1.5 times the interquartile range from the 25th and 75th percentiles; polygons represent density estimates of data and extend to extreme values.

### The PBP2B PASTA domains strengthen the interaction with DivIB

A possible explanation for the temperature sensitive phenotype of the PBP2B-ΔPASTA strain is that the PBP2B-ΔPASTA protein becomes more labile at increased temperatures. However, an analysis of PBP2B and PBP2B-ΔPASTA stability at 30°C and 48°C revealed that although PBP2B is less stable at 48°C, both PBP2B and PBP2B-ΔPASTA are degraded at similar rates (Figure S3). We noted that the temperature sensitivity of the PBP2B-ΔPASTA strain was similar to the phenotype of a *divIB* deletion strain (25). DivIB (in other organisms FtsQ) is a divisome protein that interacts with DivIC (in other organisms FtsB) and FtsL and that regulates the turnover of FtsL and DivIC (26, 27). This turnover is regulated by PBP2B and the transpeptidase domain of PBP2B has been shown to interact with the C-terminus of DivIB (27, 28). We hypothesized that the absence of the PASTA domains from PBP2B might influence the interaction with DivIB and/or other proteins. To test this, we performed a bacterial two hybrid (BACTH) assay, in which we tested the ability of PBP2B and PBP2B-ΔPASTA to interact with DivIB, DivIC, FtsL, and itself. On plate, we confirmed the previous result from Daniel and colleagues (27) that PBP2B interacts with DivIB and FtsL, but not with DivIC and found no apparent difference between PBP2B and PBP2B-ΔPASTA (Figure 4A). Notably, we did not detect a PBP2B self-interaction. Also, we only found positive results when the PBP2B variants were expressed from the pKT25 plasmid (Figure S4) – this is probably due to the difference in copy numbers between the two plasmids used in the assay and not uncommon in BACTH screens of interactions between PBPs and other proteins (29). We also analysed the interactions using a β-galactosidase assay (Figure 4B), which has the added benefit of providing a quantitative result which can give a hint about the strength of the interaction. It has to be noted that the ‘strength’ of an interaction does not scale 1:1 with β-galactosidase activity and thus that changes in activity are only indicative of a change in interaction. The β-galactosidase assay confirmed the interactions of PBP2B with DivIB and FtsL, but in the absence of the PASTA domains the activity resulting from the interaction with DivIB was roughly halved whereas the activity resulting from the interaction with FtsL was unchanged. This result suggests that the PASTA domains of PBP2B are not required for the interaction with DivIB, but that they do increase the strength of the interaction.

**Figure 4.**
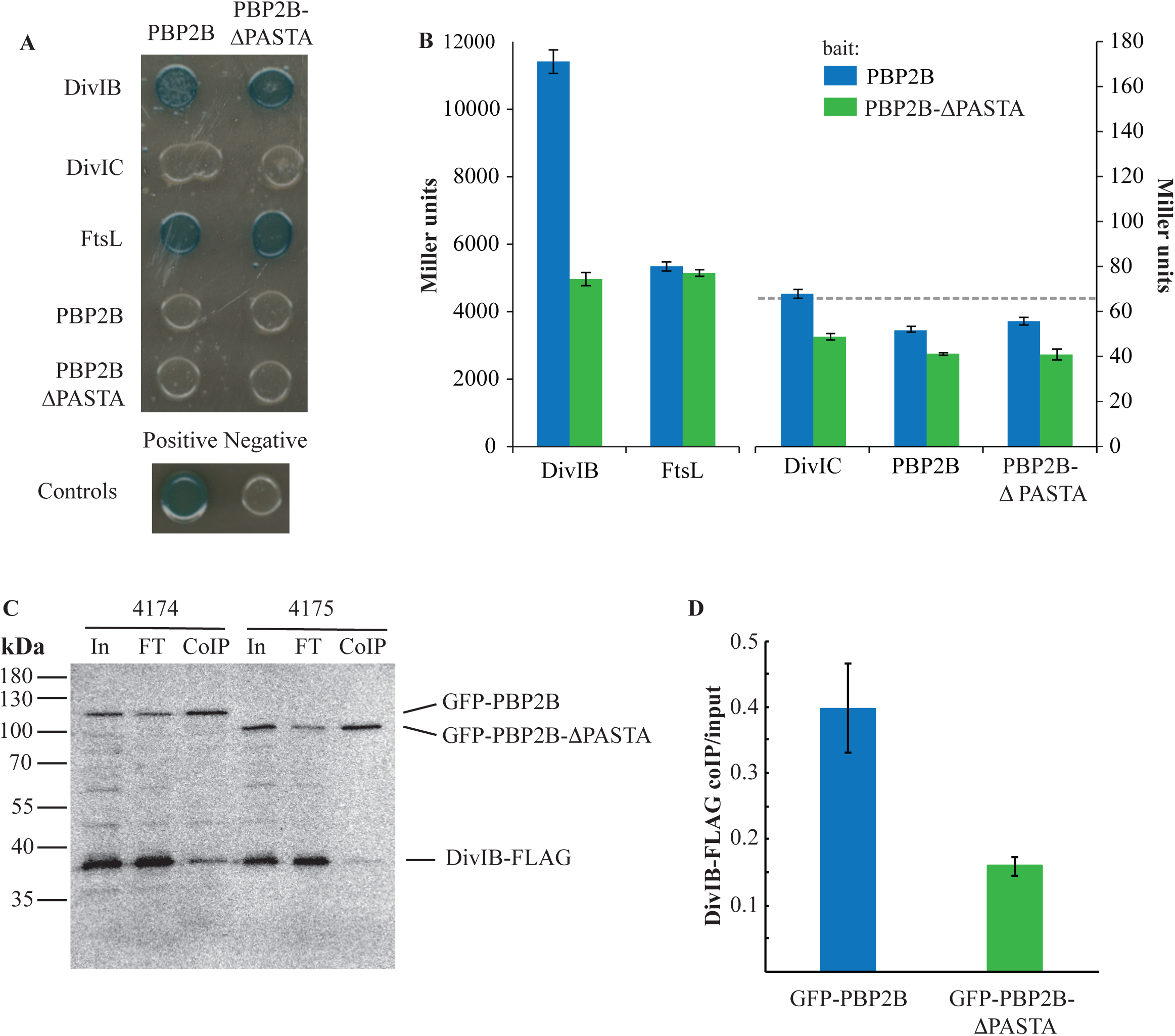
The interaction between PBP2B and DivIB is diminished in the absence of the PASTA domains. A, B) BACTH assay. A) Interaction assay on plates containing X-Gal. PBP2B, PBP2BΔ-PASTA, DivIB, DivIC and FtsL were cloned into plasmids pKT25 and pUT18C and co-transformed into *E. coli* BTH101. Co-transformants were grown on LB plates containing X-Gal and incubated at 30°C for 36 hrs. Blue colonies are considered indicative of protein-protein interaction. PBP2B and PBP2B ΔPASTA were used as bait in pKT25, while prey proteins were expressed in pUT18C. Positive control: transformants containing pKT25-zip and pUT18C-zip; negative control: transformants containing empty pKT25 and pUT18. Representative results from three independent experiments are shown. B) β-galactosidase assay. Interaction between PBP2B and PBP2B-ΔPASTA cloned into pKT25 in combination with the late division protein cloned into pUT18C. The positive control showed an activity of 63278 Miller units and the negative control 66 (shown as dotted line). Note the different scales on the left and right y-axes and the discontinued x-axis. Representative results from three independent experiments are shown. All experiments were performed in triplicates. The resulting average and standard error are shown for each interaction. C, D) Co-immunoprecipitation assay. C) Western blot of a co-Immunoprecipitation experiment. Solubilised membranes from strains producing DivIB-FLAG and GFP-PBP2B (4174) or GFP-PBP2B-ΔPASTA were immunoprecipitated using GFP-Trap agarose beads. Input (In), flow-through (FT) and eluted (coIP) material was analysed by SDS-PAGE/Western Blot and the blot was simultaneously developed using anti-FLAG and anti-GFP antibodies. D) Quantification of the fraction of DivIB-FLAG co-immunoprecipitated with either GFP-PBP2B or GFP-PBP2B-ΔPASTA, expressed as the ratio of the signal of DivIB-FLAG in the coIP fraction to the signal of DivIB-FLAG in the input sample. Bar shows the mean of 4 independent experiments with standard deviations.

To validate the results from the BACTH experiments, co-immunoprecipitation experiments were performed. GFP-PBP2B and GFP-PBP2B-ΔPASTA were produced in a *B. subtilis* strain that produces a FLAG-tagged version of DivIB at the native locus under control of the wild type promoter (GP2005, a kind gift from Jörg Stülke). DivIB-FLAG is functional as the GP2005 strain is not thermosensitive (not shown). Anti-GFP nanobodies coupled to agarose (GFP-trap) were used to immunoprecipitate GFP-PBP2B and GFP-PBPB2B-ΔPASTA and the immunoprecipitate was analysed by Western blot and detection using anti-FLAG and anti-GFP antibodies. The amount of DivIB-FLAG immunoprecipitated from cells producing GFP-PBP2B appeared significantly higher than that from cells producing GFP-PBP2B-ΔPASTA, although the overall recovery in both cases was low (Figure 4C). This was confirmed by quantification of the amount of immunoprecipitated DivIB-FLAG as a fraction of the total input in the sample (Figure 4D). Combined, the BACTH and co-immunoprecipitation results indicate that the PASTA domains of PBP2B strengthen the DivIB-PBP2B interaction.

## Concluding remarks

In this paper we show that although the PASTA domains are not absolutely essential for the scaffolding role of PBP2B, they do become essential at elevated temperatures. This phenotype is similar to the phenotype described for a *divIB* deletion (25), suggesting the PASTA domains are involved in the same pathway. We could show that the PASTA domains are involved in the interaction between PBP2B and DivIB using both a BACTH and a coimmunoprecipitation approach. Earlier, King and colleagues identified an interaction between the C-terminal part of DivIB and the transpeptidase domain of PBP2B (28). In the modelled structure of PBP2B the transpeptidase domain and the DivIB C-terminus are at similar distance from the membrane, but this is also the distance at which the PASTA domains can be found (28). It is possible that the PASTA domains strengthen the interaction between DivIB and the transpeptidase domain, but alternatively the PASTA domains interact with another region of DivIB. Our results show that PASTA domains can have distinct functions in similar proteins – whereas the PASTA domains in *S. pneumoniae* clearly function to allosterically activate the transpeptidase activity of the protein (12), their function in *B. subtilis* PBP2B is to stabilize an important protein-protein interaction in the divisome.

## Author Statements

Contributor Role

Conceptualisation DMA, DJS

Methodology DMA, AMV, LCB

Validation DMA, AMV, LCB

Formal Analysis DMA, AMV

Investigation DMA, AMV, LCB

Writing – Original Draft Preparation DMA, DJS

Writing – Review and Editing DMA, AMV, LCB, DJS

Visualisation DMA, AMV, DJS

Supervision DJS

Project Administration DJS

Funding DJS

## Acknowledgements

We would like to thank Fabian Commichau and Jörg Stülke (Göttingen University) for the kind gifts of the BACTH plasmids, and strain GP2005, respectively. We also thank Tal Shamia (Chromotek GmbH) for advice on the use of GFP-trap beads.

