## supplemental tables 1,2; supplemental figures 1-4. for "The PASTA domains of *Bacillus subtilis* PBP2B stabilize the interaction of PBP2B with DivIB"

**Figure S1. Structures of PBP2B variants used in this study.** A)PBP2B B) PBP2B-ΔPASTA C) PBP2B-SpoVD D)PBP2- PrkC. C: cytoplasmic domain, TM: transmembrane, Dim: dimerization domain, P: PASTA domain.

**Figure S2. Strains grown in SM medium.** A) Cells were grown at 30 or 37°C until exponential phase. Membranes were stained using Nile red. Representative results from three independent experiments are shown. Scale bar: 5 µm. B) Length of PBP2B strains grown on SM medium. Cells were grown in SM medium at 30 or 37°C. As *B. subtilis* forms chains, cells were labelled with Nile red in order to determine the boundaries of single cells. Length of cells was obtained by automated image analysis. The values obtained ($n = 200$ per strain) are shown as box plots. White circles show the medians (PBP2B 30°C =3.59 µm, PBP2B-ΔPASTA 30°C=4.61 µm, PBP2B 37°C =2.53 µm, PBP2B-ΔPASTA 37°C =3.45 µm); box limits indicate the 25th and 75th percentiles as determined by R software; whiskers extend 1.5 times the interquartile range from the 25th and 75th percentiles; polygons represent density estimates of data and extend to extreme values. C) Growth curves in SM medium at 48°C, $OD_{600}$ was measured every 10 min. (♦) SM medium, control, (■)168, (▲) DivIB, (●) GFP-PBP2B, (◊) GFP-PBP2B-ΔPASTA,( □) PBP2B (Δ) PBP2B-ΔPASTA. Representative results from three independent experiments are shown. All experiments were performed in triplicates. The resulting average and standard error are shown for each time point.

**Figure S3. PBP2B and PBP2B-ΔPASTA stability.** Membrane protein of GFP-PBP2B and GFP-PBP2B-ΔPASTA strains were isolated from cells incubated at 30°C. Membranes were incubated at 30°C (A, C, E) or 48°C (B, D, F) at different time points, and then incubated with bocillin. GFP was followed by fluorescence in gel (A, B) and western blot using anti-GFP antibodies (E, F). Effects on folding were followed by the ability of bocillin to bind to PBPs (C, D). The star indicates the position of GFP-PBP2B and the triangle the position of GFP-PBP2B-ΔPASTA. Representative results from three independent experiments are shown.

**Figure S4. Bacterial two-hybrid.** A) Interaction assay on plates containing X-Gal. PBP2B, PBP2BΔ-PASTA, DivIB, DivIC and FtsL were cloned into plasmids pKT25 and pUT18C and co-transformed into *E. coli* BTH101. Co-transformants were grown on LB plates containing X-Gal and incubated at 30°C for 36 hrs. Blue colonies are considered indicative of protein-protein interaction. PBP2B and PBP2B-ΔPASTA were used as bait in pUT18C, while prey proteins in pKT25. Representative results from three independent experiments are shown. B) Bacterial two hybrids β-galactosidase assay. Interaction between the fusion of PBP2B and PBP2B-ΔPASTA cloned into pUT18C in combination with the fusion of late division protein cloned into pKT25 Positive control showed an activity of 63278 Miller units and the negative control 66 (shown as dotted line). Representative results from three independent experiments are shown. All experiments were performed in triplicates. The resulting average and standard error are shown for each interaction.

**Table S1. List of primers used in this study**

| Primer | Sequence | Source |
|---|---|---|
| PrkC PASTA fw | **ACCAACTGAAAAATCTGACTCAGATAAGGAAGAA**ATGCCTAAG GATGTCAAAATACCT | This work |
| PrkC PASTA rv | **ATCGATACCGTCGACCCTCGAG**TTAGAGAGAG AATGTCACTTCA ACT | This work |
| SpoVD PASTA fw | **ACCAACTGAAAAATCTGACTCAGATAAGGAAG**G AAGACACAAAA ACAATAGAAGTTCCGA | This work |
| SpoVD PASTA rv | **ATCGATACCGTCGACCCTCGAG**TTATTCAGTCA AATACACGCGT ATC | This work |
| b2h_divIBfw | GGAGGATCTAGAGATGAACCCGGGTCAAGACC | This work |
| b2h_divIBrev | GGATTCGGTACCCCCTCAATTTTCATCTTCC | This work |
| b2h_divICfw | GGAGGATCTAGAGATGAATTTTTCCAGGGAACG A | This work |
| b2h_divICrev | GGATTCGGTACCGGCTACTTGCTCTTCTTCTCC | This work |
| b2h_ftsLfw | GGAGGATCTAGAGATGAGCAATTTAGCTTACCA ACC | This work |
| b2h_ftsLrev | GGATTCGGTACCTCATTCCTGTATGTTTTTCAC | This work |
| b2h_pbpBfw | GGAGGATCTAGAGATGATTCAAATGCCAAAAAA G | This work |
| b2h_pbpbBrev | GGATTCGGTACCTTAATCAGGATTTTAAACTTA ACCTTG | This work |
| b2h_pbpbBdP rev | GGATTCGGTACCTTATTCTTCCTTATCTGAGTCAG | This work |

* Bold letters correspond to the original plasmid sequence

**Table S2. Average of length (µm) of wild-type, PBP2B and PBP2B-ΔPASTA strains under different conditions.** Cells were grown in CH or SM medium at 30 or 37°C. As *B. subtilis* forms chains, cells were labelled with Nile red in order to determine the boundaries of single cells. Length of cells was obtained by automated image analysis. The values obtained ($n$ = 200 per strain) were averaged. The standard error is presented between brackets.

| | CH | | SM | |
|---|---|---|---|---|
| | 30°C | 37°C | 30°C | 37°C |
| wild-type (168) | 2.96 (±0.05) | 3.14 (±0.06) | 1.85(±0.03) | 1.78 (±0.04) |
| PBP2B | 3.34 (±0.06) | 3.12 (±0.05) | 3.78 (±0.09) | 2.68 (±0.83) |
| PBP2B ΔPASTA | 4.85 (±0.10) | 5.20 (±0.13) | 4.70 (±0.14) | 3.79 (±0.10) |
| PBP2B SpoVD | 3.57(±0.07) | | | |
| PBP2B PrkC | 4.55 (±0.12) | | | |

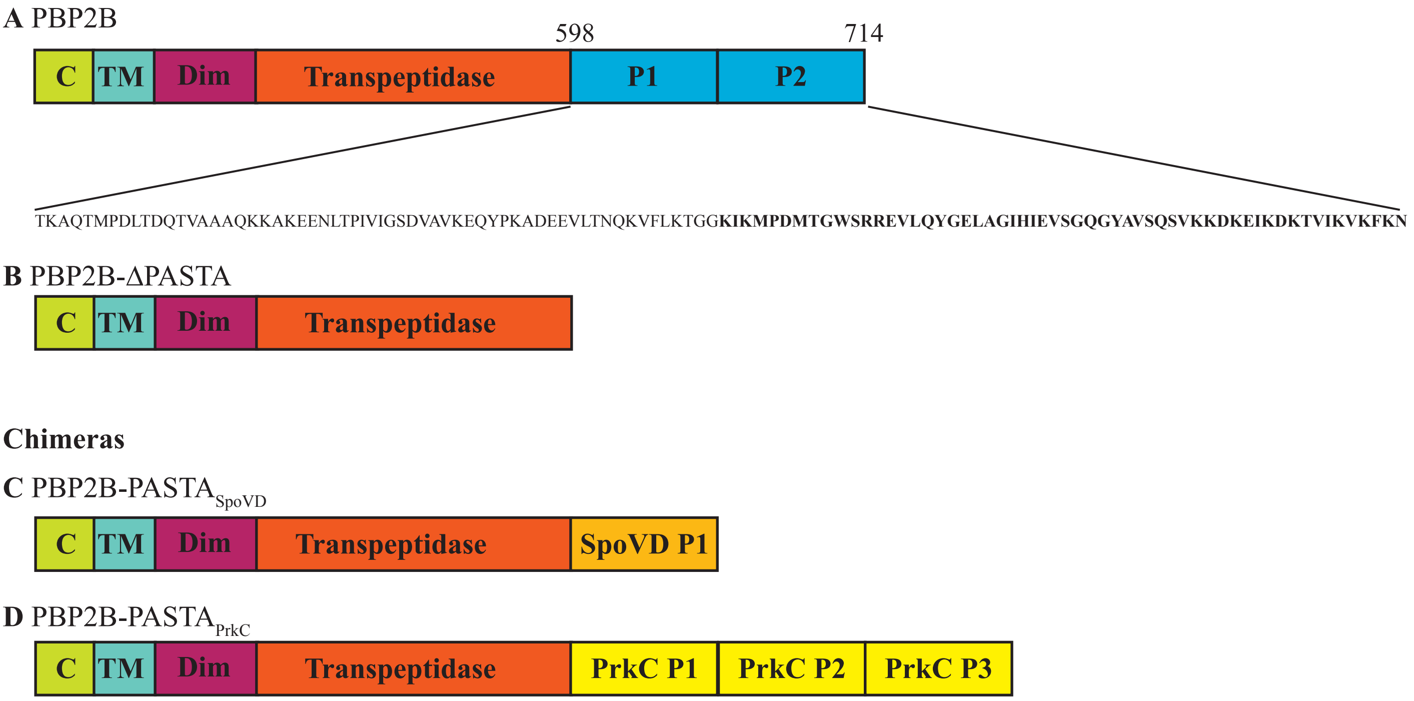
A PBP2B
598
714
C
TM
Dim
Transpeptidase
P1
P2
TKAQTMPDLTDQTVAAAQKKAKEENLTPIVIGSDVAVKEQYPKADEEVLTNQKVFLKTGG**KIKMPDMTGWSRREVLQYGELAGIHIEVSGQGYAVSQSVKKDKEIKDKTVIKVKFKN**
B PBP2B-ΔPASTA
C
TM
Dim
Transpeptidase
Chimeras
C PBP2B-PASTA SpoVD
C
TM
Dim
Transpeptidase
SpoVD P1
D PBP2B-PASTA PrkC
C
TM
Dim
Transpeptidase
PrkC P1
PrkC P2
PrkC P3

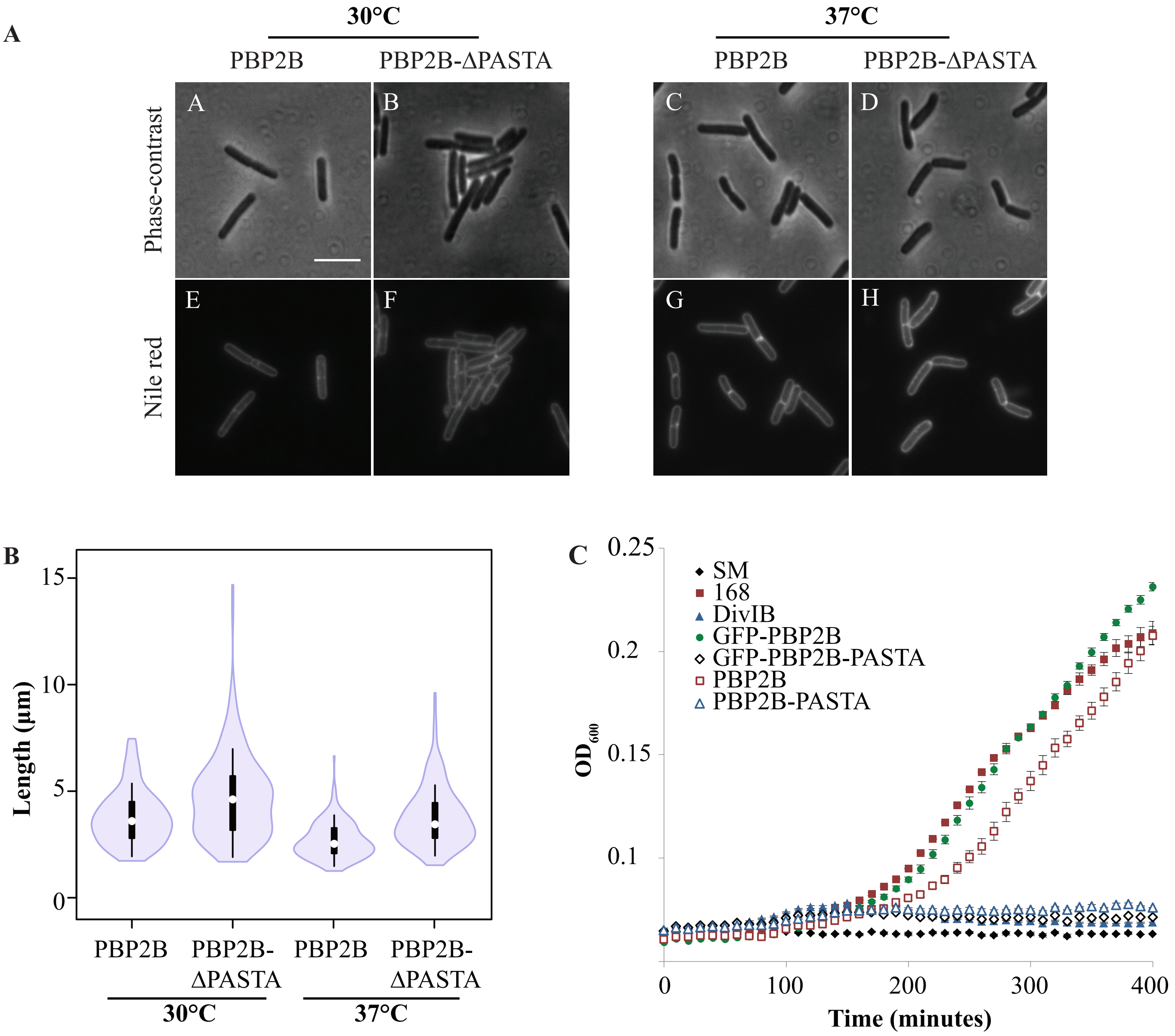
A
30°C
37°C
PBP2B
PBP2B-ΔPASTA
PBP2B
PBP2B-ΔPASTA
Phase-contrast
Nile red
A
B
C
D
E
F
G
H
B
Length (μm)
15
10
5
0
PBP2B
PBP2B-ΔPASTA
PBP2B
PBP2B-ΔPASTA
30°C
37°C
C
OD600
0.25
0.2
0.15
0.1
SM
168
DivIB
GFP-PBP2B
GFP-PBP2B-PASTA
PBP2B
PBP2B-PASTA
0
100
200
300
400
Time (minutes)

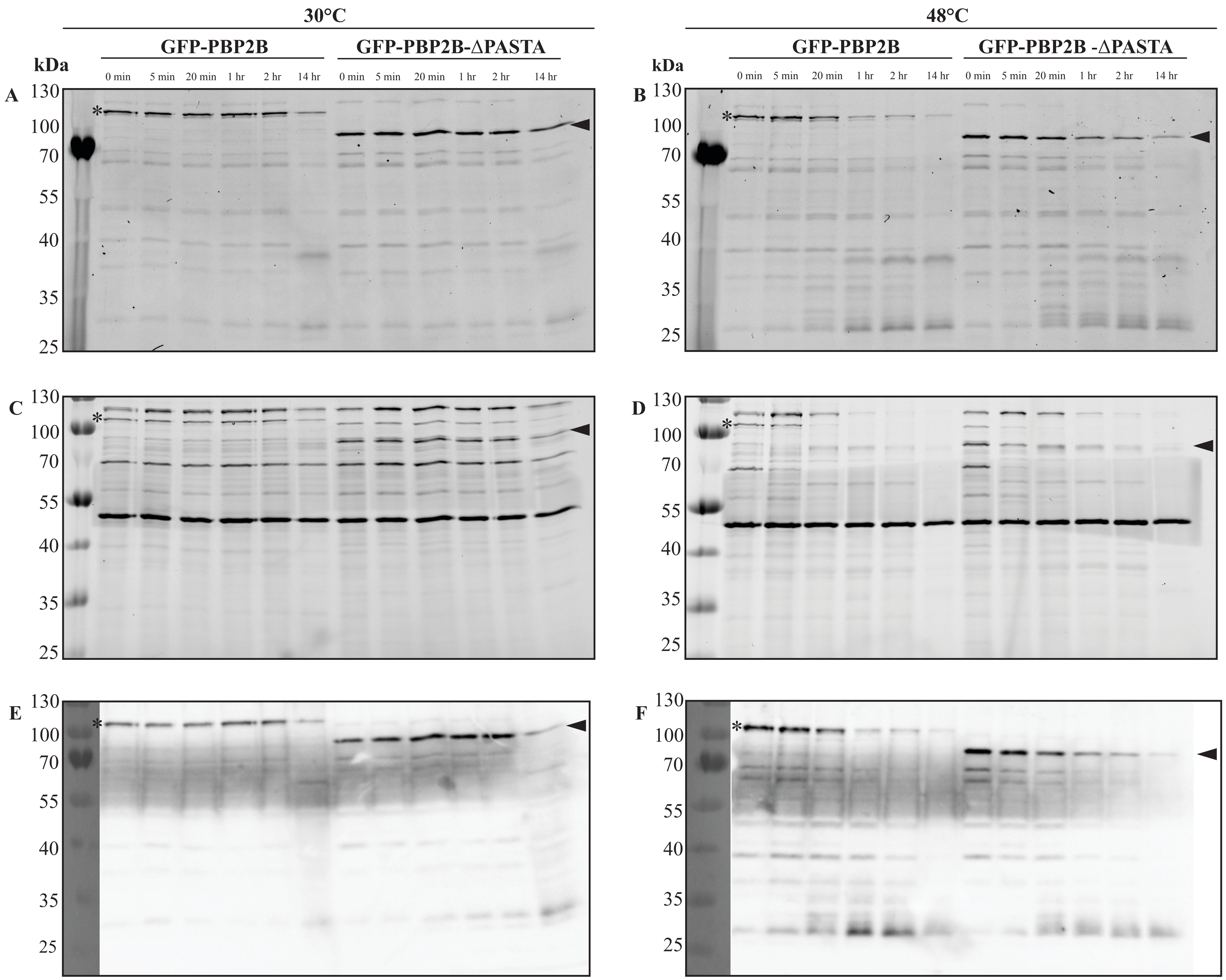

30°C
48°C
GFP-PBP2B
GFP-PBP2B-ΔPASTA
GFP-PBP2B -ΔPASTA
kDa
0 min
5 min
20 min
1 hr
2 hr
14 hr
A
B
C
D
E
F
130
100
70
55
40
35
25

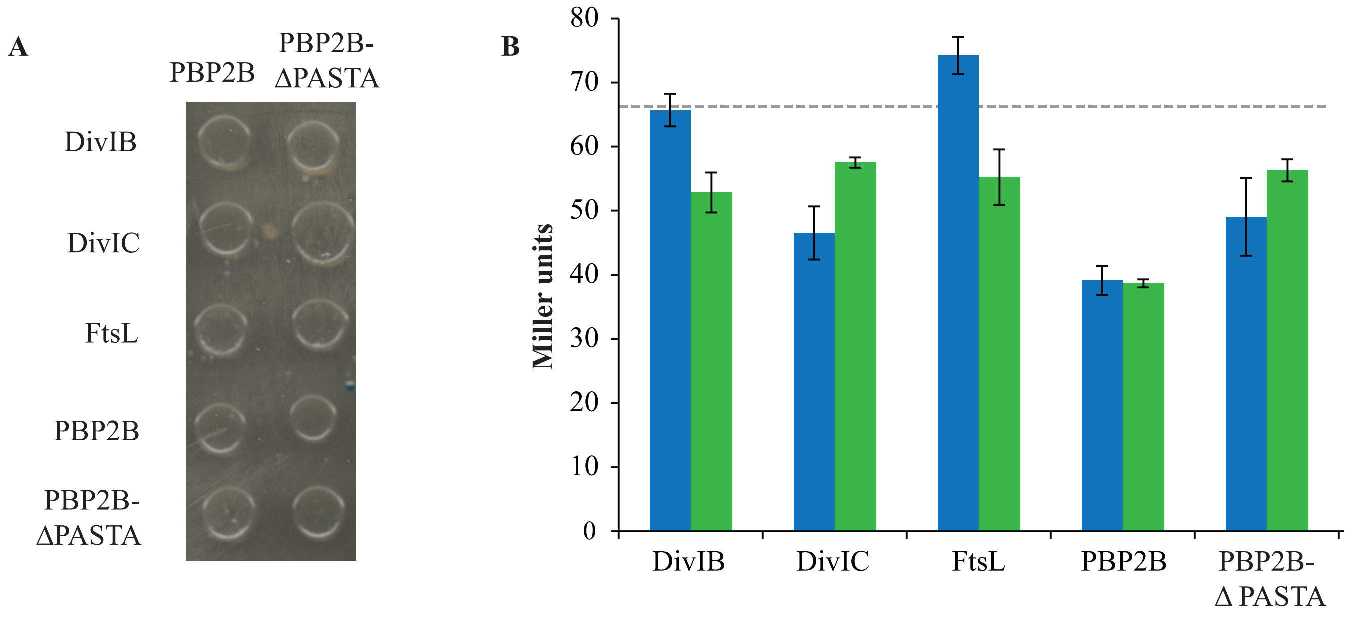

A
PBP2B
PBP2B-
ΔPASTA
DivIB
DivIC
FtsL
PBP2B
PBP2B-
ΔPASTA
B
Miller units
80
70
60
50
40
30
20
10
0
DivIB
DivIC
FtsL
PBP2B
PBP2B-
Δ PASTA
